# Microthrombocytopenia caused by impaired microtubule stability in RhoB-deficient mice

**DOI:** 10.1101/2021.11.04.467272

**Authors:** Maximilian Englert, Katja Aurbach, Annika Gerber, Tobias Heib, Isabelle C. Becker, Lou M. Wackerbarth, Charly Kusch, Ayesha A. Baig, Sebastian Dütting, Ulla G. Knaus, Christian Stigloher, Bernhard Nieswandt, Irina Pleines, Zoltan Nagy

**Affiliations:** Institute of Experimental Biomedicine, University Hospital, University of Würzburg, Würzburg, Germany; Rudolf Virchow Center for Integrative and Translational Bioimaging, University of Würzburg, Würzburg, Germany; Conway Institute, School of Medicine, University College Dublin, Dublin, Ireland; Imaging Core Facility, Biocenter, University of Würzburg, Germany

## Abstract

Megakaryocytes are large cells in the bone marrow, which give rise to blood platelets. Platelet biogenesis involves megakaryocyte maturation, the localization of mature cells in close proximity to bone marrow sinusoids and the formation of protrusions, which are shed into the circulation. Rho GTPases play important roles in platelet biogenesis and function. RhoA-deficient mice display macrothrombocytopenia and a striking mislocalization of megakaryocytes into bone marrow sinusoids and a specific defect in G-protein signaling in platelets. However, the role of the closely related protein RhoB in megakaryocytes or platelets remains unknown. In this study, we show that, in contrast to RhoA deficiency, genetic ablation of RhoB in mice results in microthrombocytopenia (decreased platelet count and size). RhoB-deficient platelets displayed mild functional defects predominantly upon induction of the collagen/glycoprotein VI pathway. Megakaryocyte maturation and localization within the bone marrow, as well as actin dynamics were not affected in the absence of RhoB. However, *in vitro* generated proplatelets revealed pronouncedly impaired microtubule organization. Furthermore, RhoB-deficient platelets and megakaryocytes displayed selective defects in microtubule dynamics/stability, correlating with pronouncedly reduced levels of acetylated α-tubulin. Our findings imply that absence of this tubulin posttranslational modification results in decreased microtubule stability leading to microthrombocytopenia in RhoB-deficient mice. Our data thus points to specifically impaired microtubule - but not actin - dynamics as a general mechanism underlying the manifestation of microthrombocytopenia *in vivo*. We furthermore demonstrate that RhoA and RhoB have specific, non-redundant functions in the megakaryocyte lineage.

**KEY POINTS:**

- RhoB-deficient mice display microthrombocytopenia
- RhoB has different functions in the megakaryocyte lineage than RhoA and regulates microtubule dynamics

## INTRODUCTION

Blood platelets are small anucleated cells with central roles in hemostasis and thrombosis. Platelets derive from large precursor cells called megakaryocytes (MKs) that are found predominantly in the bone marrow (BM). MK development is characterized by endomitosis (DNA replication without cell division) and cytoplasmic maturation, which comprises the biosynthesis of platelet-specific granules and the formation of an internal demarcation membrane system (DMS), which serves as a membrane reservoir for the produced platelets. Platelet biogenesis is a complex cellular process,^1,2^ initiated by mature MKs in direct contact with the BM sinusoids,^3^ and involves the penetration of endothelial cells,^4,5^ and the extension of large cytoplasmic protrusions into the sinusoidal lumen,^6,7^ which are shed and undergo maturation in the peripheral circulation to become *bona fide* platelets. The different steps of platelet biogenesis highly depend on rearrangements of both the microtubule (MT) and actin cytoskeletal network.^1,2,8^ Consistently, genetic defects impairing cytoskeletal dynamics can result in thrombocytopenia in humans and mice.^9^

Rho GTPases act as molecular switches by cycling between an inactive guanosine diphosphate (GDP)-bound and an active guanosine triphosphate (GTP)-bound state. In the latter conformation, Rho GTPases interact with multiple effector proteins to regulate diverse processes including cytoskeletal dynamics, cell adhesion, cell migration, cell polarity, cell cycle and vesicle trafficking. Multiple members of the Rho family have been established as critical regulators of platelet production and function.^10–12^ We have previously reported that cytoplasmic MK maturation and subsequent transendothelial platelet biogenesis requires balanced signaling of RhoA and Cdc42.^13,14^ Mice lacking RhoA in the MK lineage display macrothrombocytopenia (decreased platelet count and increased platelet size), accompanied by a pronounced mislocalization of whole MKs inside of BM sinusoids.^14,15^ Higher vertebrates express three Rho subfamily proteins, RhoA, RhoB and RhoC, which are 87% homologous.^16^ In immune cells, RhoB is involved in cytokine trafficking and cell survival,^17^ affects cell adhesion and migration through β2 and β3 integrins,^18^ but is not required for podosome assembly in macrophages.^19^ A variety of lipid modifications within RhoB promotes its localization to the plasma membrane, endosomes and multivesicular bodies.^20^

In this study, we investigated for the first time the role of RhoB in platelet biogenesis and function by using constitutive knockout mice. We show that, in contrast to RhoA deficiency, loss of RhoB results in *micro*thrombocytopenia (decreased platelet count and size), which was accompanied by moderately impaired platelet function. Our results demonstrate that RhoA and RhoB have specific, independent functions in the MK lineage and point to selective defects in MT dynamics in RhoB-deficient mice as a mechanism underlying the manifestation of microthrombocytopenia *in vivo*.

## METHODS

### Mice

Constitutive RhoB-deficient mice (further referred to as RhoB^-/-^)^21^ were kindly provided by Ulla G Knaus. Animal studies were approved by the district government of Lower Franconia (Bezirksregierung Unterfranken).

### Platelet count and size by flow cytometry

Platelet count and size were determined by flow cytometry with size determined beads using a FACSCalibur (BD Biosciences), as previously described.^22^

### Proplatelet formation of MKs

Proplatelet formation of BM-derived *in vitro*-differentiated MKs was analyzed 72 h after MK isolation following the lineage depletion protocol. MKs were seeded into 96-well-plates and incubated for 48 h and counted at different time points manually at a Zeiss PrimoVert brightfield microscope. Brightfield images were acquired at an Evos Microscope (ThermoFisher Scientific). For fluorescent visualization of the cytoskeleton, coverslips were coated with 0.1% Poly-L-Lysine for 20 min at RT and washed with H_2_O, before 500 μl of 1x platelet buffer and 300 μl of the cell suspension were gently added to the wells. By centrifugation for 4 min at 900 rpm proplatelets were forced down on the coverslip. Cells were fixed with 800 μl 4% PFA in PHEM containing 0.1% TritonX100 and then blocked in 3% BSA for 1 h. MKs were stained with anti-α-tubulin Alexa488 (3.33 mg/ml) and phalloidin-Atto647N (170 nM), and analyzed by confocal microscopy at a Leica TCS SP8. Organization of the microtubule network was assessed by focusing on thickness and shape of proplatelet shafts and tips compared to the WT controls.

### Statistics

Data presented are mean ± standard deviation (SD). Unpaired, two-tailed t-test or Mann-Whitney test were used to determine statistical significance between two groups with normal or non-normal distribution, respectively. Sample size (n) and statistical significance are reported in the figures or figure legends. Stars indicate statistically significant differences compared to *wt* (* p ≤ 0.05, ** p ≤ 0.01 and *** p ≤0.001). Statistical analysis was performed in GraphPad Prism 9 (GraphPad Software).

Platelet life span, integrin activation, degranulation, glycoprotein expression, aggregation, adhesion under flow conditions, spreading on fibrinogen, cold-induced microtubule disassembly, tail bleeding time, intravital thrombosis model, immunofluorescence microscopy, cold-induced microtubule disassembly, dSTORM, TEM analysis, MK isolation, spreading, and immunoblotting are described in detail in the Supplemental Methods.

## RESULTS

### *RhoB^-/-^* mice display microthrombocytopenia

The role of the RhoB in platelets and MKs was studied in constitutive *RhoB^-/-^* mice^21^ and compared to wild-type (*wt*) littermate controls. The complete loss of RhoB on the protein level in platelets from *RhoB^-/-^* mice was confirmed by immunoblotting (**Figure 1A**). The protein levels of other important Rho GTPases, namely RhoA, Cdc42 and Rac1 remained unaffected in the absence of RhoB, indicating no direct compensatory regulation among these proteins.

**Figure 1.**
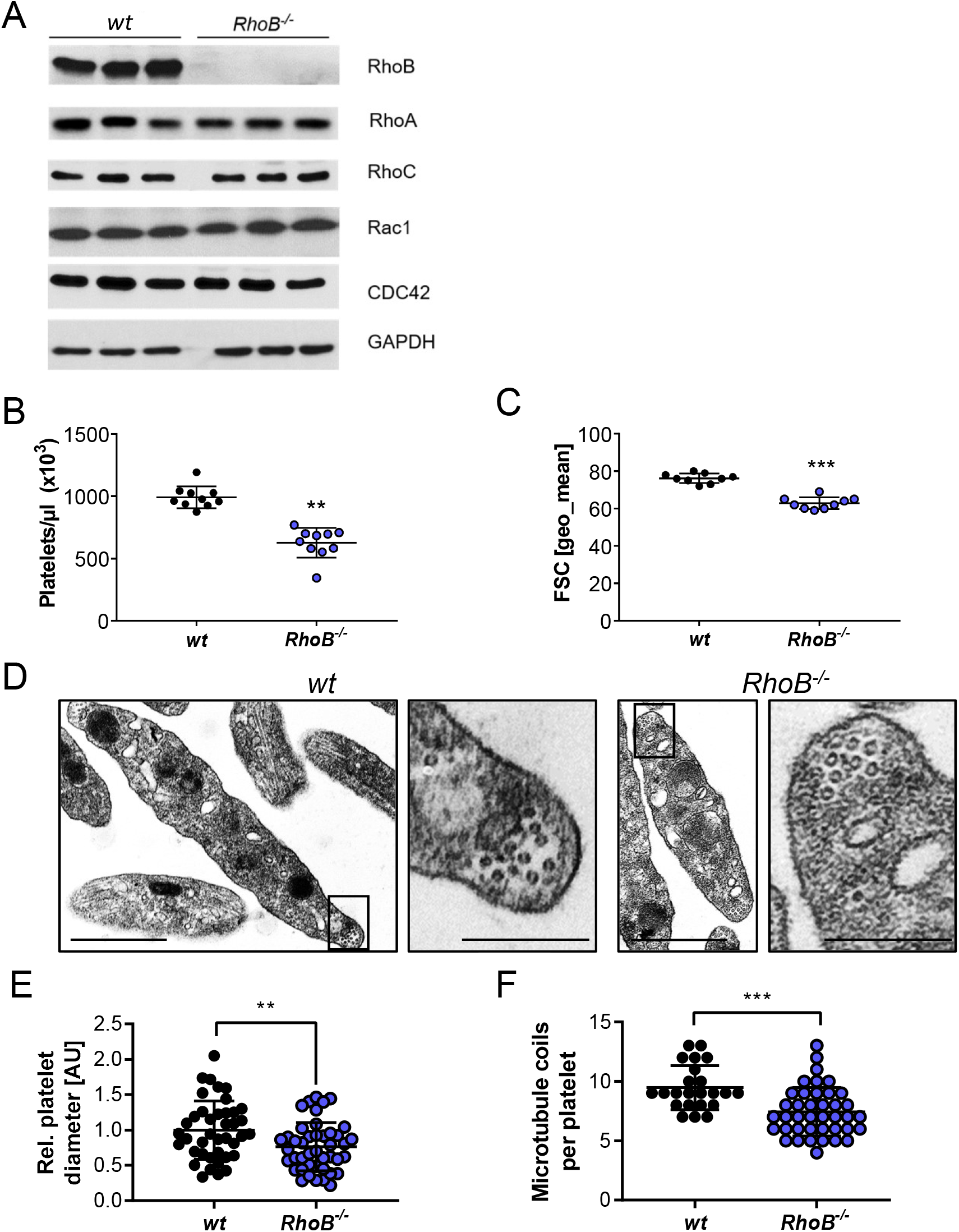
Loss of RhoB leads to microthrombocytopenia and alters microtubule organization in circulating platelets. (A) Blots of platelet lysates from wt and *RhoB^-/-^* mice immunoblotted for the indicated proteins. GAPDH was used as loading control (n=3). (B) Platelet count assessed by flow cytometry. Each data point represents one mouse (n=12). (C) Platelet size assessed by forward scatter characteristics determined by flow cytometry. Each data point represents one mouse (n=12). (D) Representative images of ultrastructural analysis of resting *wt* (left) and *RhoB^-/-^* (right) platelets by transmission electron microscopy (TEM). Scale bar overview: 1 μm; inset: 0.5 μm. (E) Quantification of platelet diameter (relation of platelet width to length) of *RhoB^-/-^* platelets relative to *wt* using TEM images described in D (*wt* n=41; *RhoB^-/-^* n=43). (F) Quantification of microtubule (MT) coils/platelet using TEM images described in D. MT number was determined by manual count of at least 5 images/genotype. Each data point represents one single platelet (*wt* n=23; *RhoB^-/-^* n=40). **p < 0.01; ***p < 0.001; Mann-Whitney test, mean ± SD.

While RhoA-deficient mice display a characteristic macrothrombocytopenia,^15^ loss of RhoB led to microthrombocytopenia (**Figure 1B-C; supplemental Table 1**). Other peripheral blood parameters were largely unaltered indicating a specific role of RhoB in the MK lineage. The decreased platelet size of *RhoB^-/-^* platelets was confirmed by transmission electron microscopy (TEM) analysis (**Figure 1D-E**). Resting *wt* platelets contain an average of 8 – 12 MT coils, which are located in the periphery to maintain their spherical shape. Notably, the number of MT coils in *RhoB^-/-^* platelets was significantly reduced (*wt*: 9.48 ± 1.86 vs. *RhoB^-/-^* 7.43 ± 2.05 MT coils per platelet) (**Figure 1D, F**). The distribution of α- and dense (δ-) granules, on the other hand, was not affected by loss of RhoB (**supplemental Figure 1A-B**). These results demonstrate that RhoB regulates platelet count and size *in vivo*.

### Moderately impaired platelet function in *RhoB^-/-^* mice

We next determined the characteristics and function of *RhoB^-/-^* platelets *in vitro* by flow cytometry. Levels of surface glycoproteins (GPs) were not significantly different between *RhoB^-/-^* and *wt* platelets, except for a slight reduction in GPV and the collagen receptor GPVI (**supplemental Table 2**). Platelet activation induces a shape change from a resting, discoid to a spherical form.^23^ RhoA-deficient platelets show selective defects in platelet activation and shape change after stimulation of agonists that induce in G-protein-coupled (Gα_13_, Gα_q_) signaling pathways, which result in impaired intravital thrombosis and a mild bleeding phenotype.^15^ To investigate whether absence of RhoB had a similar effect, we analyzed platelet shape change and compared to RhoA-deficient platelets by light transmission aggregometry upon stimulation with a low dose of the stable thromboxane A2 analog U46619 (**Figure 2A**). Shape change in *RhoB^-/-^* platelets was unaltered, while it was abrogated in RhoA-deficient platelets, demonstrating that RhoB is not involved in Gα_13_ signaling. However, the aggregation of *RhoB^-/-^* platelets in response to intermediate concentrations of thrombin, U46619 or convulxin (CVX) was moderately reduced indicating an overall impairment of platelet function (**Figure 2B**).

**Figure 2.**
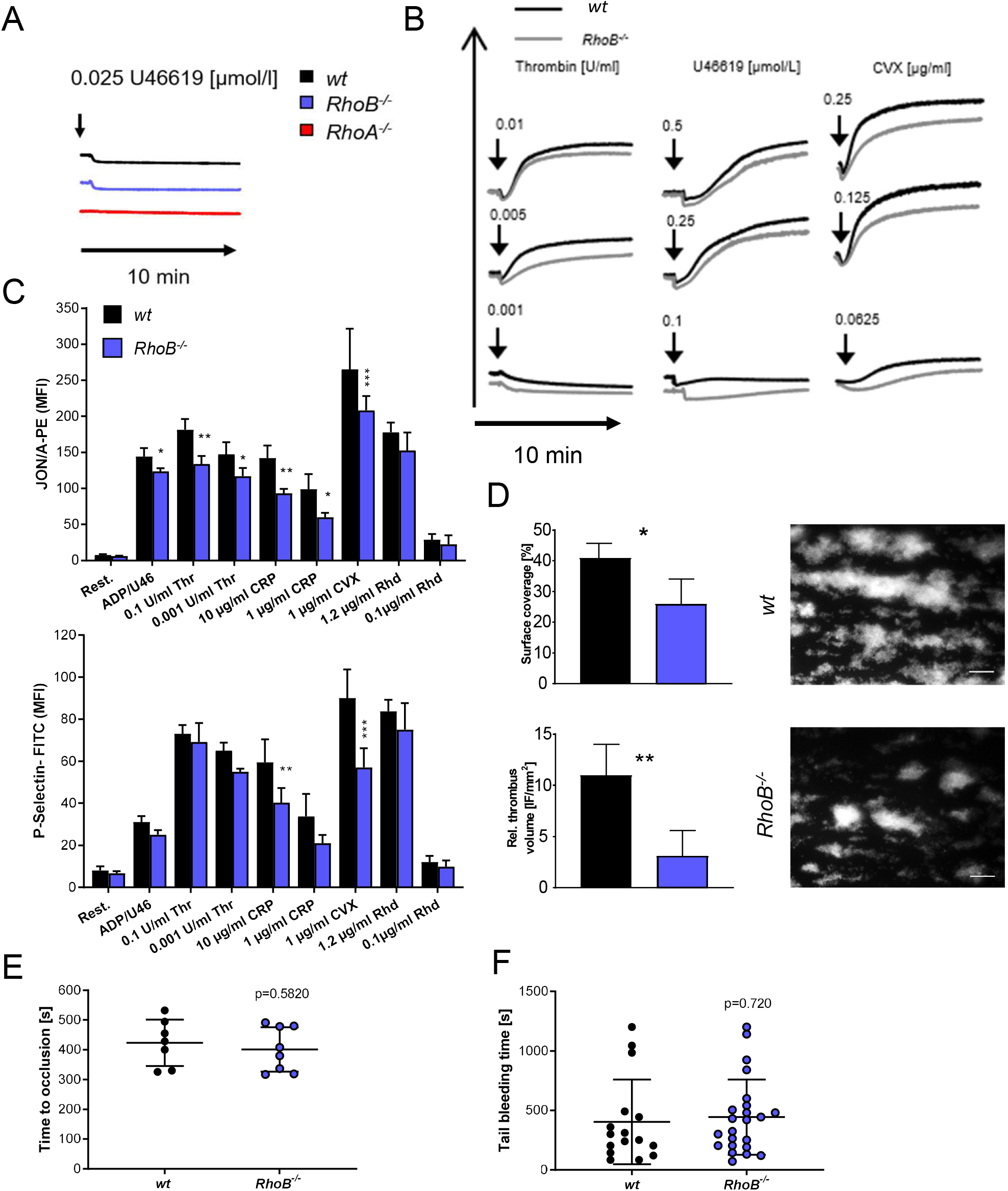
Impaired activation and thrombus formation of *RhoB^-/-^* platelets *in vitro*. (A-B) Platelet light transmission aggregometry. Representative aggregation curves of washed *wt* and *RhoB^-/-^* platelets stimulated with different agonist concentrations under stirring conditions for 10 min are shown. (A) Shape change of *wt* and *RhoB^-/-^* platelets in response to low concentrations of the thromboxane analog U46619. RhoA-deficient platelets displaying defective shape change under these conditions were used as a positive control. (B) Aggregation of *RhoB^-/-^* platelets *in vitro*. (C) Platelet integrin αIIbβ3 activation (JON/A-PE binding; upper) and α-granule secretion (assessed by P-Selectin surface exposure; lower) *in vitro* were determined upon stimulation of *wt* and *RhoB^-/-^* platelets with the indicated agonists at different concentrations (n=4). (D) *Ex vivo* thrombus formation under flow on a collagen-coated surface. Representative images of thrombus formation at an intermediate shear of 1000 s^-1^ are shown. Upper panel: Adhesion of platelets to collagen assessed by area coverage. Lower panel: Formation of 3D aggregates displayed as relative thrombus formation (n=8). (E) Induction of arterial thrombosis by mechanical injury of the abdominal aorta of *wt* and *RhoB^-/-^* mice *in vivo*. Each data point represents one mouse (*wt* n=7; *RhoB^-/-^* n=8). (F) Tail bleeding time on filter paper of *wt* and *RhoB^-/-^* mice *in vivo*. Each data point represents one mouse (*wt* n=15; *RhoB^-/-^* n=23; data were pooled from 3 independent experiments). *p < 0.05; **p < 0.01; ***p < 0.001; Mann-Whitney test, mean ± SD. Rest = resting; ADP/U466 = Adenosine diphosphate/U46619; Thr = thrombin; CRP = collagen-related peptide; CVX = convulxin; Rhd = Rhodocytin.

To investigate this finding further, we next assessed platelet activation by flow cytometry, which allows for a more detailed analysis of the activation response. In contrast to the reported selective defect of RhoA-deficient platelets in response to thrombin and U46619,^15^ *RhoB^-/-^* platelets displayed moderately reduced activation in response to most tested agonists. The most profound defect was observed upon stimulation of the collagen/GPVI signaling pathway with either collagen-related peptide (CRP) or CVX (**Figure 2C**). This was consistent for both integrin αIIbβ3 activation (as assessed by JON/A-PE binding) as well as degranulation (as assessed by binding of anti-P-Selectin-FITC).

It was previously demonstrated that *RhoB^-/-^* macrophages displayed reduced level of β2 and β3, but not β1 integrins.^19^ Since murine platelets do not express β2 integrins and the surface expression of β3 and β1 integrins was unaltered in *RhoB^-/-^* platelets (**supplemental Table 2**), we investigated platelet β1 integrin activation (as assessed by 9EG7-FITC binding) by flow cytometry and found similar activation defects as for αIIbβ3 integrin activation (**supplemental Figure 1C**).

In order to more closely mimic *in vivo* thrombus formation, we next performed *ex vivo* flow chamber assays, in which anticoagulated whole blood was perfused over a collagen-coated surface at intermediate shear rate (1000 s^-1^). *RhoB^-/-^* platelets were able to form structured thrombi under these conditions (**Figure 2D**), however, the overall surface coverage and relative thrombus volume were significantly reduced compared to *wt* platelets. This was observed for both intermediate and particularly at high shear rates and, in accordance with our *in vitro* results, indicated that loss of RhoB predominantly affected the GPVI pathway. The observed platelet function defects *in vitro* did not translate into altered arterial thrombus formation *in vivo*, which was determined by mechanical injury of the abdominal aorta (**Figure 2E**). Furthermore, unaltered tail bleeding times indicated that hemostasis was not affected in *RhoB^-/-^* mice (**Figure 2F**). These findings stand in contrast to the defects observed in RhoA-deficient mice *in vivo* and emphasize that RhoA and RhoB are involved in different platelet signaling pathways.

### Unaltered platelet turnover in *RhoB^-/-^* mice

Decreased circulating platelet counts may manifest as a result of impaired platelet generation, an increased platelet clearance rate, or a combination of both. We have previously reported that macrothrombocytopenia in RhoA-deficient mice is associated with increased platelet turnover.^15^ We therefore investigated the lifespan of circulating, anti-GPIb-labeled platelets over a period of five days by flow cytometry (**Figure 3A**). We observed only a minor reduction of the platelet lifespan in *RhoB^-/-^* mice, indicating that the thrombocytopenia was predominantly caused by a direct impairment of platelet production. *RhoB^-/-^* mice exhibited splenomegaly, however the overall spleen morphology and splenic MK numbers were unaltered (**Figure 3B-D**). The presence of splenomegaly might be a consequence of the lack of RhoB in other hematopoietic cells. Platelet removal was reported to be modulated by desialylation of surface glycoproteins, which are recognized by Ashwell-Morell receptors and, together with *Macrophage galactose lectin* (MGL), mediate clearance of platelets by hepatocytes.^24,25^ However, *RhoB^-/-^* platelets did not show alterations in *Erythrina cristagalli lectin* (ECL) binding to exposed galactose (**Figure 3E**) and in terminal galactose levels (determined by the ratio of neuraminidase treated platelets versus untreated platelets) (**Figure 3F**). Moreover, the level of RNA-rich, *thiazole orange* (TO)-positive platelets, an indicator of young, reticulated platelets, was not altered (**Figure 3G**). Taken together, these results indicate that the thrombocytopenia in *RhoB^-/-^* mice was not caused by increased platelet clearance, but by a defect in either MK maturation or defective platelet generation.

**Figure 3.**
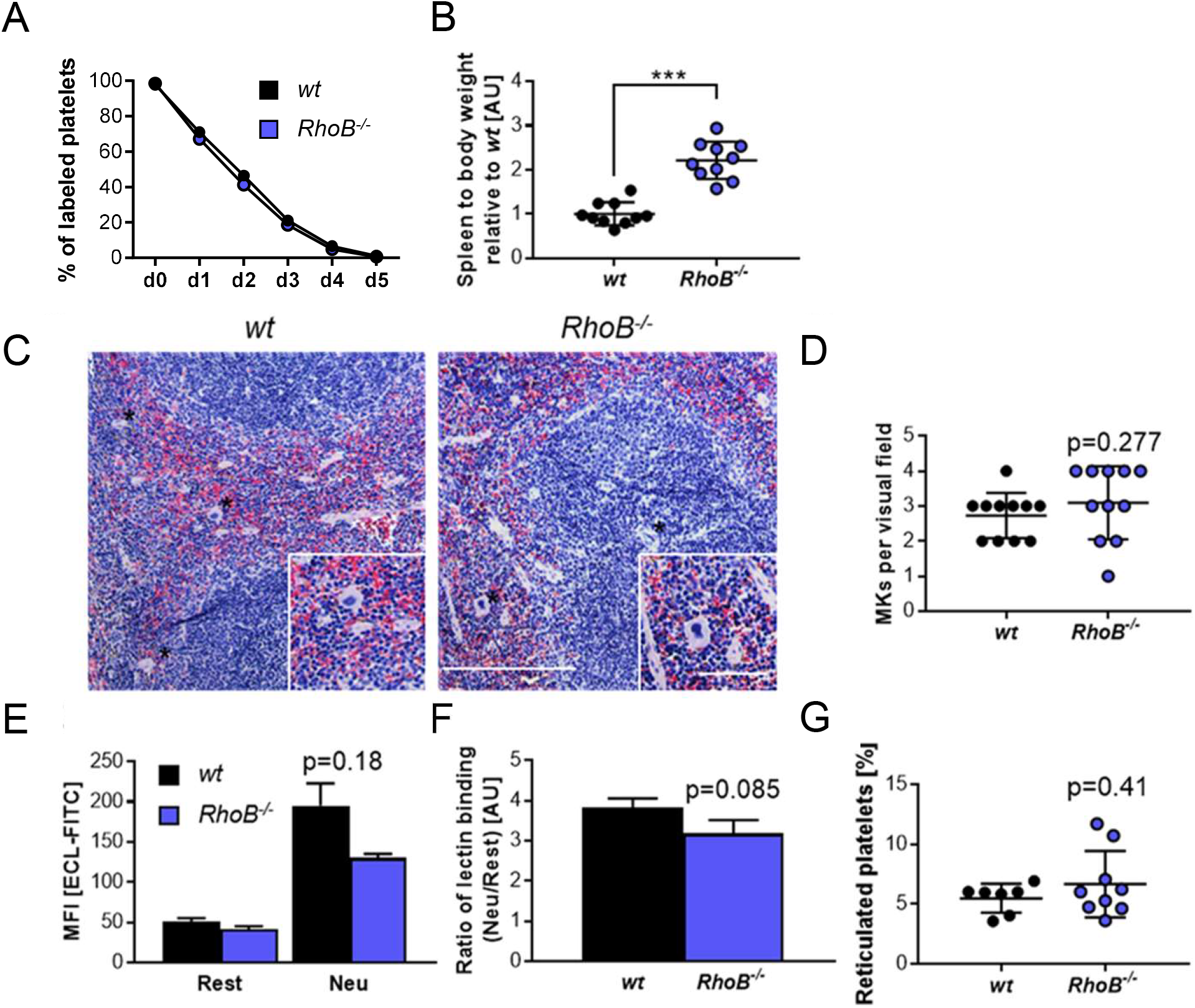
Lack of increased platelet turnover in *RhoB^-/-^* mice. (A) For the determination of platelet lifespan, *wt* and *RhoB^-/-^* platelets were labeled with an α-GPIX antibody and the labeled platelet population was analyzed over five consecutive days (n=5 mice/genotype). *p < 0.05; **p < 0.01; ***p < 0.001; multiple unpaired 2-tailed Student’s t test with Holm-Sidak correction for multiple comparisons, mean ± SD (B) Ratio of spleen weight (mg) to body weight (mg) of *RhoB^-/-^* mice relative to *wt* (n=10 mice/genotype). (C) Paraffin sections of murine spleens stained using hematoxylin and eosin (H&E). Analysis was performed at a Leica DMI4000 B microscope. Scale overview: 100 μm, inset: 80 μm. (D) Number of MKs/visual field determined in spleen paraffin sections (n=3; 4 visual fields/genotype). (E) Assessment of ECL-FITC binding to *wt* and *RhoB^-/-^* platelets by flow cytometry at resting state (rest) or upon neuraminidase treatment (Neu) (n=3). (F) Ratio of lectin binding (neuraminidase treated platelets versus untreated platelets) for both genotypes (n=3). (G) Percentage of reticulated (RNA-rich) platelets assessed by TO binding in flow cytometry (*wt* n=7; *RhoB^-/-^* n=9). ***p < 0.001; Mann-Whitney test, mean ± SD.

### Impaired microtubule organization and proplatelet formation in *RhoB^-/-^* MKs

Given the pivotal role of Rho GTPases, in the modulation of cytoskeletal dynamics, we next investigated the outcome of RhoB deficiency on MK maturation and function. MK numbers in the BM compartment were unaltered (**Figure 4A**). Investigation of the ultrastructure of BM MKs by TEM *in situ* revealed a normally developed DMS and granules in *RhoB^-/-^* MKs (**Figure 4B**). *In situ* analysis of BM cryosections stained for CD105 (vessel marker) and GPIX (MK marker) revealed that RhoB had no influence on MK localization (**Figure 4C-D**). Together, these results indicated that RhoB is dispensable for MK maturation and localization *in vivo*. This is in marked contrast to the intrasinusoidal MK mislocalization observed in RhoA-deficient mice, and thus emphasized differential functions of RhoB and RhoA not only in platelets, but also MKs.

**Figure 4.**
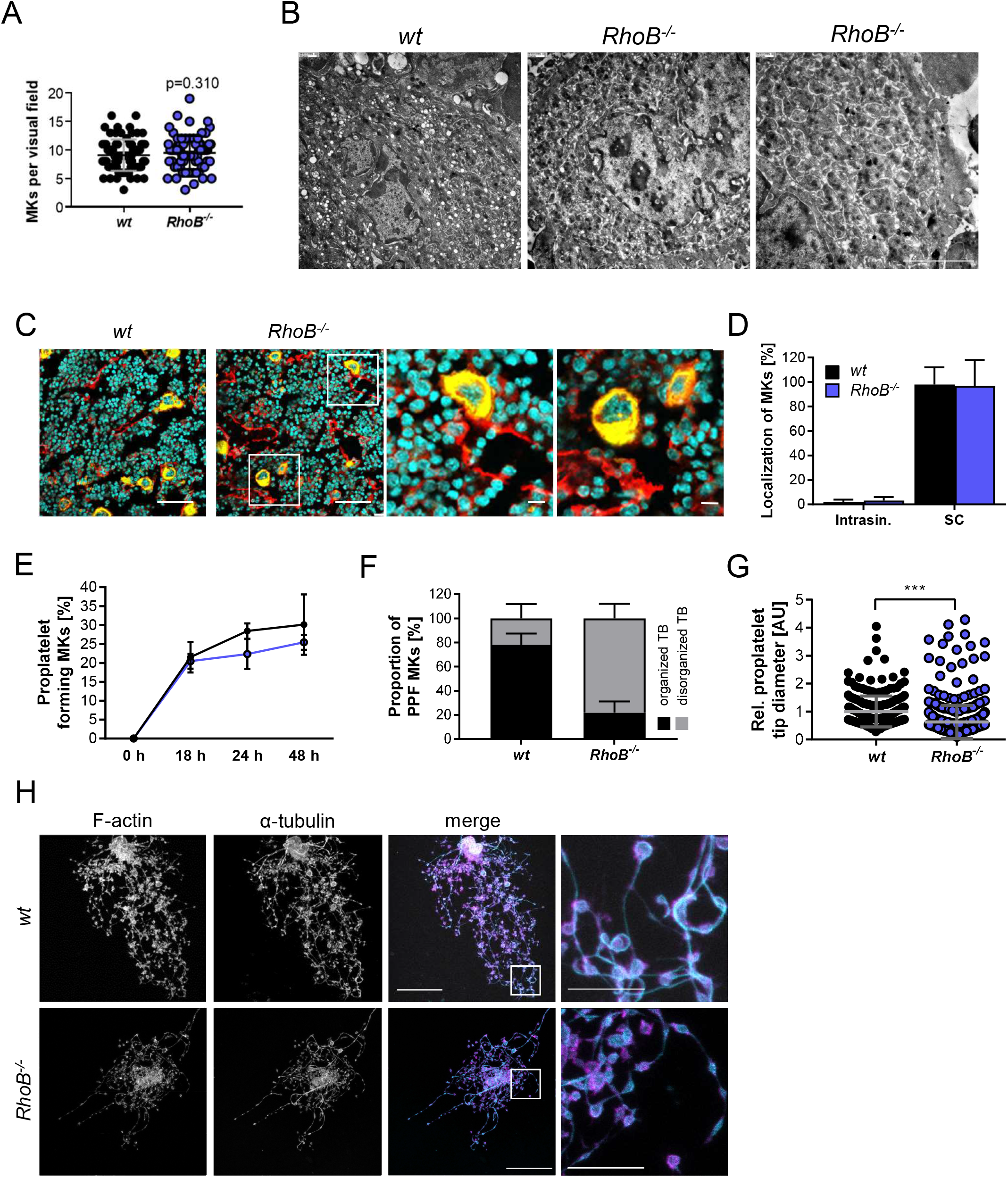
Decreased proplatelet tip size and impaired microtubule organization of *RhoB^-/-^* MK *in vitro*. (A) Quantification of MK numbers in *wt* and *RhoB^-/-^* BM. At least 10 images/mouse were analyzed using 7 mice/genotype. Each data point represents one MK (n =75). Unpaired, two-tailed t-test, mean ± SD. (B) Representative images of ultrastructural analysis of *wt* and *RhoB^-/-^* MKs *in situ* using TEM. Scale bar: 4 μm. (C) Representative confocal microscopy images of femur cryosections. The femora were processed using the Kawamoto-method for cryosections. Sections were stained with fluorescently-labeled anti-CD105 antibody to stain vessels, and fluorescently-labeled anti-GPIX antibody to visualize MKs and DAPI (nucleus). Scale bar in overview equals 40 μm and in zoom 5 μm. (D) Quantification of MK localization in *wt* and *RhoB^-/-^* BM in images shown in B. (E-H) Analysis of proplatelet formation of BM-derived *RhoB^-/-^* MKs. (E) Quantification of *in vitro* proplatelet formation at 18 h, 24 h and 48 h after enrichment by BSA gradient (n=3 mice/genotype). Multiple unpaired 2-tailed Student’s t test with Holm-Sidak correction for multiple comparisons, mean ± SD. (F) Quantification of microtubule organization in enriched *wt* and *RhoB^-/-^* MKs (n=3 mice/genotype). At least 13 images/mouse were analyzed. Mann-Whitney test, mean ± SD. (G) Quantification of proplatelet tip diameter of *wt* and *RhoB^-/-^* MKs 24 h after BSA gradient (normalized to *wt*, n=2 mice/genotype). At least 11 MKs/mice were analyzed. Each data point represents one measured proplatelet tip (*wt* = 250; *RhoB^-/-^* = 494). ***p < 0.001; Mann-Whitney test, mean ± SD. (H) Representative confocal images of *in vitro*-differentiated MKs 24 h after BSA gradient. The MKs were spun down to Poly L-Lysine coated glass slides and stained with phalloidin-Atto647 (magenta), α-tubulin-Alexa488 (cyan) and DAPI to visualize the nucleus. Scale bar in overview: 40 μm; inset: 5 μm. Intrasin. = intrasinusoidal; SC = sinusoidal contact; TB = tubulin.

We next analyzed the ability of *RhoB^-/-^* MKs to produce proplatelets *in vitro*. For this, BM-derived MKs were cultured for 3 days in the presence of TPO and hirudin to induce proplatelet formation,^26^ followed by staining for F-actin and α-tubulin (**Figure 4E-H**). We could not detect alterations in the kinetics of proplatelet formation between enriched *wt* and *RhoB^-/-^* MKs during the observation time of 48 h (**Figure 4E**), indicating that the ability to fragment into proplatelets was still preserved in the absence of RhoB. Furthermore, the distribution of F-actin appeared unaffected in *RhoB^-/-^* proplatelets. Strikingly, a high proportion (78.1%) of proplatelets formed by *RhoB^-/-^* MKs displayed a disorganized microtubule (MT) network, as evident by the enlarged proplatelet shafts and irregular-shaped proplatelet tips with an overall decreased size (**Figure 4F-H; supplemental Figure 2A**).

Since the analysis of proplatelets provides only limited information on the functionality of Factin dynamics in the absence of RhoB, we additionally analyzed the characteristic integrin- and actin-dependent formation of podosomes in *in-vitro*-differentiated MKs upon adhesion on collagen-coated cover slips. Importantly, podosome formation along the collagen fibers, as well as the overall signal intensity of F-actin was similar in *wt* and *RhoB^-/-^* MKs (**supplemental Figure 2B-C, E**), Consistently, and in contrast to observations of MKs with impaired podosome formation, e.g. upon Cdc42 or Profilin1 deficiency,^13,27^ loss of RhoB did not affect the total MK spreading area (**supplemental Figure 2D**). The distribution of tubulin was also not obviously altered between wt and *RhoB^-/-^* MKs under these adhesive conditions (**supplemental Figure 2F**) pointing to a specific requirement of RhoB in the regulation of MT dynamics under the conditions of proplatelet formation. Collectively, these results demonstrated that RhoB is a critical modulator of MT, but not actin dynamics in MKs during the process of proplatelet formation, which might provide an explanation for the decreased size of circulating *RhoB^-/-^* platelets *in vivo*.

### RhoB is a critical regulator of microtubule stability in MKs and platelets

To further investigate the mechanism underlying the tubulin organization defect during proplatelet formation upon loss of RhoB, we investigated important regulators of the MK cytoskeleton in MKs. Total protein levels of α-tubulin and β1-tubulin (exclusively expressed in the MK lineage) were unaltered BM-derived *in vitro*-differentiated MKs (**supplemental Figure 3A**). Protein levels of non-muscle myosin IIa (NMIIa) and IIB (NMIIb), that are downstream effectors of the Rho subfamily and influence cell migration and contractility, were also unaltered. Furthermore, in line with functional podosome formation in *RhoB^-/-^* MKs, levels of the podosome-associated proteins vinculin and Arp2 were not affected by RhoB deficiency (**supplemental Figure 3B**). The levels of the MT plus-end tracking protein *Adenopolyposis Coli* (APC), which was recently reported as a negative regulator of proplatelet formation in mice,^28^ was also unchanged in *RhoB^-/-^* MKs (**supplemental Figure 3A**).

These findings indicated that RhoB may directly influence tubulin dynamics during platelet formation. Due to the heterogeneity of MK cultures, we decided to focus on the circulating platelets, i.e. the terminal *in vivo* products of proplatelet formation in our further investigations. We analyzed MT and F-actin distribution in adherent, spread platelets *in vitro*. For this, platelets were incubated for 5, 15 and 30 min on fibrinogen-coated cover slips, which induces extensive spreading of *wt* platelets that is supposed to be driven by integrin outside-in signaling. Similar to the normal adhesion and spreading of MKs on collagen, we did not observe a difference in the spreading kinetics between *wt* and *RhoB^-/-^* platelets (**Figure 5A-B**) indicating that integrin outside-in signaling is functional in the absence of RhoB.

**Figure 5.**
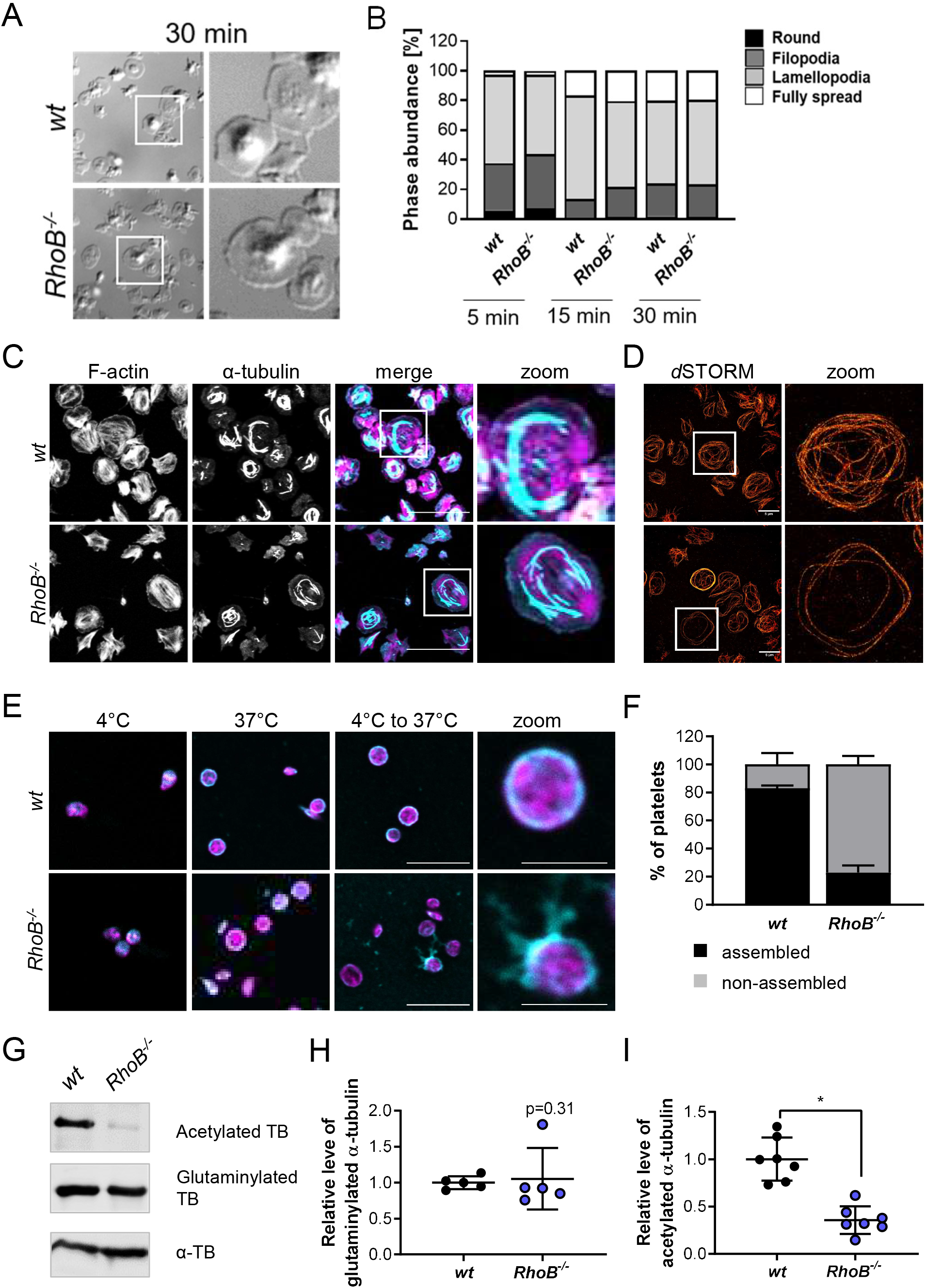
Impaired microtubule stability in *RhoB^-/-^* platelets. (A-C) Spreading and F-actin dynamics in *RhoB^-/-^* platelets. (A) Representative images of platelets spread on fibrinogen-coated coverslips for 30 min. (B) Quantification of platelet spreading on fibrinogen for 5, 10 and 15 min (n=3). *p < 0.05; Mann-Whitney test, mean ± SD. (C-D) Analysis of spread platelets on fibrinogen by immunofluorescence confocal and superresolution (dSTORM) microscopy. (C) Representative confocal microscopy images of platelets spread on fibrinogen for 30 min with 0.1 U/ml thrombin, stained with phalloidin-Atto647 (magenta) and α-tubulin-Alexa488 (cyan) and imaged using a Leica TC SP8 confocal microscope. Scale bar in overview: 10 μm. (D) Representative dSTORM images of platelets spread on fibrinogen for 30 min, stained with α-tubulin-Alexa488 (red) and imaged using an in house made dSTORM microscope. Scale bar: 5 μm. (E-F) Analysis of cold-induced MT disassembly and re-assembly of spread platelets *in vitro*. (E) Representative confocal immunofluorescence microscopy images of MT organization at 4°C, 37°C and a combination of disassembly at 4°C with following reassembly at 37°C in platelets. Platelets were stained with phalloidin-Atto647 (magenta) and α-tubulin-Alexa488 (cyan). Scale bar in overview: 10 μm; inset: 3 μm. (F) Quantification of microtubule organization in *wt* and *RhoB^-/-^* platelets after cold-induced MT disassembly and reassembly (4°C to 37°C). At least 5 images/mouse were analyzed (n=3). *p < 0.05; Mann-Whitney test, mean ± SD. (G-I) Analysis of acetylation and detyrosination (glutamylation) of platelet α-tubulin residues by immunoblotting. (G) Platelet lysates of *wt* and *RhoB^-/-^* mice were immunoblotted for post-translational modifications (PTM) of α-tubulin (acetylated α-tubulin and glutaminylated α-tubulin). (H-I) Quantification of PTM immunoblots. Each data point represents one mouse (acetylated α-tubulin n=7; glutaminylated α-tubulin n=5). *p < 0.05; Mann-Whitney test, mean ± SD.

To analyze the distribution of F-actin and MT, spread platelets were stained for α-tubulin and F-actin. In line with results from MKs, the distribution of F-actin was not affected by RhoB deficiency. Consistently, F-actin content and assembly upon CRP or thrombin stimulation were unaltered in *RhoB^-/-^* platelets (**supplemental Figure 3D**), emphasizing that RhoB is dispensable for F-actin dynamics in MKs and platelets.

In contrast, the α-tubulin network appeared markedly altered in *RhoB^-/-^* platelets with more shortened MTs (**Figure 5C**). We therefore next visualized the MT network in spread platelets by superresolution microscopy (*d*STORM), as described.^29^ Indeed, we observed a high proportion of *RhoB^-/-^* platelets with few peripheral MT coils and an overall looser MT network compared to the *wt* (**Figure 5D**). These results were in line with the decreased number of tubulin coils in resting *RhoB^-/-^* platelets (**Figure 1D, F**) and pointed to a critical role of RhoB in MT stabilization.

MTs are highly dynamic structures that are constantly assembled and disassembled. MT disassembly into dimers can be artificially induced by exposure to cold (4°C) and reassembly into fibers occurs spontaneously at 37°C.^30^ To investigate the role of RhoB in MT dynamics and stability, we induced MT disassembly in platelets by cold storage (4°C) for 2 h followed by a reassembly at 37°C.^31^ While both *wt* and *RhoB^-/-^* platelets displayed MT coils at their periphery at 37°C which disassembled upon cold-storage (**Figure 5E-F**), only *RhoB^-/-^* platelets were not able to reassemble MT coils after disassembly (wt: 83 ± 2% vs. *RhoB^-/-^:* 23 ± 5 %) (**Figure 5F; supplemental Figure 3E**).

MTs can be modified through post-translational modifications (PTMs), which serve for the adaption to different cellular functions.^32^ In platelets, PTMs of α-tubulin have been reported to be involved in MT rearrangements, including acetylation of residue K40 and detyrosination/tyrosination (in C-terminal tyrosines, often also called glutaminylation).^33^ Acetylation and detyrosination are indicators for stable, longer-lived MTs, while more dynamic MTs are tyrosinated and deacetylated.^32^ We therefore investigated the acetylation status of the α-tubulin K40 residue by immunoblotting in platelet lysates. *RhoB^-/-^* platelets showed pronouncedly decreased levels of acetylated α-tubulin (Figure 5G-H), which was by tendency also observed in MK lysates (supplemental Figure 3C). In contrast, levels of detyrosinated (glutaminylated) α-tubulin were unaltered in both platelets and MKs (Figure 5G, I; supplemental Figure 3C). Thus, deficiency in RhoB results in short-lived MT that are either *per se* more unstable or constantly remodeled. In summary, these results reveal that RhoB is a pivotal regulator of MT turnover and stability, and strongly indicate that the impaired MT dynamics is the underlying reason for defective platelet biogenesis and microthrombocytopenia in *RhoB^-/-^* mice.

## DISCUSSION

Here, we show that, in contrast to MK-specific deficiency of RhoA or Cdc42,^15,34^ loss of the small Rho GTPase RhoB results in microthrombocytopenia in mice, which is caused by a profound defect in MT stability.

The majority of platelet disorders associated with mutations in cytoskeletal genes are characterized by macrothrombocytopenia, which is reflected in transgenic mouse models of the respective diseases.^9^ Microthrombocytopenia, on the other hand, is a rare clinical condition. It is a characteristic of the *Wiskott-Aldrich Syndrome* (WAS), and X-linked thrombocytopenia (XLT), both of which are caused by mutations in the WAS gene,^35,36^ as well as CARST, a congenital auto-recessive disease caused by mutations in Adhesion and Degranulation-promoting Adaptor Protein (ADAP).^37,38^ Both WAS protein (WASP) and ADAP are primarily known as regulators of the actin cytoskeleton: ADAP is a scaffolding protein involved in receptor-induced actin cytoskeletal dynamics in platelets.^29^ WASP is a downstream effector of Cdc42 and, via the Arp2/3 complex, leads to actin cytoskeletal rearrangements associated with filopodia formation. In line with this pathway, loss of Arp2 in MKs in mice,^39^ or mutations in the Arp2/3 complex component *ARPC1B* in humans leads to microthrombocytopenia.^40^ Interestingly, deficiency of Profilin-1, another protein associated with actin turnover, in MKs results in a WAS-like phenotype in mice.^27^

Profilin-1 and Arp2 deficiency is linked to altered MT stability,^27,39^ which resembles our observations on the *RhoB^-/-^* mice. However, there are two striking differences: First, a major characteristic of mouse models with WAS- or CARST-like phenotypes is the undirected release of proplatelets into the BM compartment, referred to as “ectopic platelet release”.^27,29,36,39^ Second, platelets and MKs lacking ADAP, Arp2 or WASP display pronounced defects in Factin dynamics, including aberrant podosome formation. In contrast, *RhoB^-/-^* MKs do not show signs of ectopic platelet release and our results indicate largely normal F-actin organization, including podosome formation *in vitro*. Furthermore, protein levels of Profilin-1 and Arp2 are unaffected in the absence of RhoB. Together, our findings clearly point to impaired MT - but not actin - dynamics as the major mechanism underlying the manifestation of microthrombocytopenia *in vivo*. Our results imply that the characteristic phenotype of ectopic platelet release is critically dependent on aberrant actin dynamics upon dysregulation of the WASP/Arp2/3 pathway.

In this study, we provide evidence that in MKs and platelets RhoB and RhoA have different functions. RhoA-deficient mice display largely normal MK maturation, however transmigration of entire MKs into BM sinusoids results in macrothrombocytopenia.^15^ The phenotypic difference might be explained by the unique subcellular localization of RhoB compared to RhoA.^17^ Furthermore, in contrast to the selective Gα_13_/Gα_q_ signaling defect in RhoA-deficient platelets, *RhoB^-/-^* platelets showed an impaired platelet activation *in vitro* independent of the agonist used, but most pronounced upon stimulation of the collagen receptor GPVI. Interestingly, RhoB activity was shown to regulate Rac1 translocation to endosomes in endothelial cells.^41^ Since Rac1 is a critical regulator of PLCγ2 activation downstream of GPVI in platelets,^42,43^ loss of RhoB may affect Rac1 localization and function in platelets causing predominantly GPVI-related signaling defects. However, in contrast to Rac1-deficiency, lamellipodia formation was not affected by absence of RhoB.

RhoB has mostly been described as a regulator of actin dynamics in diverse cell types where it, similar to RhoA, regulates stress fiber formation,^44^ and cell migration.^17,20^ Interestingly, the F-actin distribution of *RhoB^-/-^* platelets and MKs appeared unaltered, which may indicate that RhoA is able to compensate for the loss of RhoB in regulating actin dynamics in the MK lineage. In contrast, *RhoB^-/-^* platelets displayed a pronounced MT assembly defect, evident by the inability to reassemble tubulin coils after cold-storage, altered α-tubulin distribution in spread platelets and decreased numbers of MT coils in resting platelets. MT dynamics are critical for the formation of proplatelets during the later stages of thrombopoiesis *in vitro* and *in vivo*. Consistently, and in line with our observations in platelets, the MT organization of *RhoB^-/-^* proplatelets was profoundly impaired, resulting in aberrantly sized and overall smaller proplatelet tips. These results thus provide a direct explanation for the decreased size of circulating platelets in *RhoB^-/-^* mice. This stands in contrast to the phenotype of *Tubb1^-/-^* mice, in which the complete loss of tubulin β1 results in a pronounced macrothrombocytopenia.^45^ These findings imply that altered tubulin regulation can have different consequences compared to tubulin deficiency.

Mechanistically, our results indicate that the marked inability to acetylate α-tubulin, a posttranslational modification associated with stable, long-lived MTs leads to decreased MT stability in *RhoB^-/-^* platelets and MKs and, subsequently, the manifestation of microthrombocytopenia. Previous studies on the platelet marginal MT band have shown that α-tubulin is heavily acetylated in platelets and contributes to the kinetics of platelet spreading.^46,47^ Furthermore, acetylation levels of K40 of α-tubulin are associated with increased tubulin longevity^32^ suggesting that decreased acetylation levels might correspond to either direct MT instability or increased MT turnover. This may explain the shortened MT coils in spread *RhoB^-/-^* platelets. Moreover, the finding that *RhoB^-/-^* platelets are not able to rebuild MT coils after cold-induced disassembly suggests that the formed MT coils are more susceptible to stress.

Regarding signaling pathways that may link RhoB to MT dynamics, the formin mDia and RhoB were shown to interact in endosomes by control of actin dynamics that were necessary for vesicle trafficking.^48^ Moreover, mice deficient for both mDia and RhoB had abnormally shaped erythrocytes, splenomegaly, and extramedullary hematopoiesis, indicating overlapping roles for RhoB and mDia in hematopoietic cells.^49^ As mDia is a downstream effector of small Rho GTPases and induces actomyosin contractility by NMIIa activation, we investigated NMIIa and NMIIb expression in *RhoB^-/-^* platelets and MKs, but did not detect a difference compared to *wt*. Loss of the MT plus end tracking protein APC in MKs and platelets resulted in decreased α-tubulin acetylation at K40.^28^ However, protein levels of APC were unaltered in *RhoB^-/-^* MKs. Additional target proteins linking RhoB to the MT cytoskeleton could be *twinfilin 1* (twf1) and cofilin1, which were recently shown as modulators of the actin/tubulin crosstalk during platelet biogenesis.^50^ Together, these findings indicate that RhoB may directly regulate tubulin acetylation, and thereby MT stability.

In summary, our results reveal that impaired MT stability, in combination with normal cytoplasmic MK maturation (including DMS formation) causes microthrombocytopenia in *RhoB^-/-^* mice *in vivo*. While genetic RhoB deficiency in humans has not been described to date, mutations resulting in persistent RhoB activation were shown to result in systemic capillary leak syndrome,^51^ and an increased risk of cerebral palsy.^52^ The analysis of platelets from these patients may provide new insights into the role of RhoB in the MK lineage in humans.

## ACKNOWLEDGEMENTS

We thank Stefanie Hartmann, Sylvia Hengst, Birgit Midloch and Daniela Naumann for excellent technical assistance and the microscopy platform of the Bioimaging Center Würzburg for providing technical infrastructure and support. This work was funded by the Deutsche Forschungsgemeinschaft (DFG, German Research Foundation) (project number 374031971 - TRR 240/project A01 to IP and BN). ZN was supported by a grant of the German Excellence Initiative to the Graduate School of Life Sciences (GSLS), University of Würzburg.

## AUTHORSHIP CONTRIBUTIONS

Conceptualization: ME, KA, BN, IP, ZN; Investigation: ME, KA, AG, TH, ICB, LMW, CK, AAB, SD, IP; Resources: UGK, CS, BN, IP; Supervision: BN, IP, ZN; Writing - original draft: ME, KA, BN, IP, ZN; Writing—review & editing: UGK, ME, BN, IP, ZN

