## Supplemental Material for "Microthrombocytopenia caused by impaired microtubule stability in RhoB-deficient mice"

#### SUPPLEMENTAL METHODS

##### Platelet lifespan

Anesthetized mice were injected i.v. with 5 µg of a Dylight 488 conjugated anti-GPIX antibody derivative. Labeled platelets were measured at the indicated timepoints by flow cytometry of diluted whole blood using a FACSCalibur.

##### Platelet integrin activation and degranulation

Anesthetized mice were retro-orbitally bled up to 50 µl into 300 µl heparin (20 U/ml in TBS), the blood washed twice in Tyrode´s buffer without $Ca^{2+}$ and centrifuged for 5 min at 2800 rpm and RT. It was then carefully resuspended in 1 ml Tyrode´s buffer with $Ca^{2+}$ for platelet activation. 50 µl of washed blood were incubated with a 1:1 mixture of antibodies directed against αIIbβ3 integrin (JON/A-PE, Emfret Analytics) and the α-granule specific protein P-selectin (WUG 1.9-FITC, Emfret Analytics) or integrin β1 (9EG7-FITC). Platelets were activated by addition of agonists of GPCRs (thrombin or the combination of the thromboxane A2 analog U46619 and ADP), or the (hem)ITAM receptors GPVI (collagen-related peptide (CRP) or convulxin (CVX)) or CLEC-2 (rhodocytin) for 6 min at 37°C. The reaction was stopped by the addition of 500 µl PBS and the mean fluorescence intensity (MFI) of the respective fluorophore coupled antibodies was assessed by flow cytometry using a FACSCalibur.

##### Platelet glycoprotein expression

Anesthetized mice were retro-orbitally bled up to 50 µl into 300 µl heparin (20 U/ml in TBS) and the blood was further diluted in 600 µl PBS. 50 µl of the diluted blood were incubated with 10 µl of fluorophore-conjugated antibodies for 20 min at RT and stopped with 500 µl PBS. The GP surface expression was subsequently determined on resting platelets by assessing the MFI of indicated fluorophore-conjugated antibodies using a FACSCalibur.

##### Platelet aggregation

Anesthetized mice were retro-orbitally bled up to 50 µl into 300 µl heparin (20 U/ml in TBS) and platelets were isolated from whole blood as previously described,[24] and adjusted to 500,000 platelets/µl before an incubation of 30 min at 37°C. 1.5 x $10^6$ platelets were suspended in Tyrode´s containing $Ca^{2+}$ and 100 µg/ml fibrinogen in cuvettes and activated by 100-fold concentrated agonists of GPCRs (thrombin, or the thromboxane A2 analog U46619) or the ITAM receptor GPVI (CVX). Thrombin stimulation was performed without the addition of fibrinogen. Light transmission was assessed over time (600 s) using a four-channel aggregometer (APACT, Laborgeräte und Analysensysteme).

##### Platelet adhesion under flow conditions

24 x 50 mm coverslips were coated with 70 µg/ml HORM collagen at 37°C o/n and specific bindings were blocked with 1% BSA in PBS. Mice were bled to 1 ml into 300 µl heparin [20

U/ml in TBS], the whole blood was diluted 1:2 in Tyrode´s buffer supplemented with $Ca^{2+}$ and the platelets were incubated with a Dylight-488-coupled anti-GPIX antibody derivate [0.2 µg/ml] for 5 min at 37°C. The coated coverslips were covered with a transparent flow chamber with a slit depth of 50 µm. The blood was then perfused over the coverslip for 4 min using a pulse free pump and washed with Tyrode's buffer containing 2 mM $Ca^{2+}$ for 4 min at the same shear rate. Brightfield (BF) and fluorescent images were taken by a Zeiss HBO 100 (Axiovert, 200M, Zeiss) and thrombus volume and surface coverage were analyzed using ImageJ Software.

**Tail bleeding time (filter paper method)**

Mice were anesthetized with triple narcotics (Midazolam [5 µg/g], Medetomidine [0.5 µg/g], Fentanyl [0.05 µg/g]) and their hemostasis was analyzed. 2 mm of the tail tip were removed with a scalpel and the forming drop of blood was gently absorbed with a filter paper every 20 sec without touching the wound until the bleeding stopped or stopped manually after 20 min by cauterization to prevent excessive blood loss.

**Intravital thrombosis model**

*In vivo* thrombus formation was performed as previously described.[1] The abdominal cavity of anaesthetized mice was opened to expose the abdominal aorta. An ultrasonic flow probe (0.5PSB699; Transonic Systems) was placed around the abdominal aorta, and thrombus formation was induced by a single firm compression with a forceps upstream of the flow probe. Blood flow was monitored for 30 min or until vessel occlusion occurred (blood flow stopped for > 5 min).

**Cold-induced microtubule disassembly**

Platelets were washed and adjusted to 500,000 platelets/µl and diluted 1:2 with Tyrode´s buffer without $Ca^{2+}$ in a final volume of 50 µl per sample or condition. Samples were either incubated at 4°C for 2 h for MT depolymerization and subsequent reassembly was allowed at 37°C for 30 min, or the platelets were kept at 37°C or at 4°C for 30 min. After incubation, the platelets were fixed and permeabilized in PHEM buffer supplemented with 1.5% PFA and 0.075% IGEPAL CA-630 and allowed to adhere to a poly-L-lysine-coated coverslip. Subsequently the platelet cytoskeleton was visualized, using Phalloidin-Atto647N and anti-α-tubulin-Alexa488 antibodies for 1 h at RT and analyzed by confocal microscopy at a Leica TCS SP5 or SP8 using a 100x objective.

**Platelet spreading on fibrinogen**

Coverslips (24 x 60 mm) were coated with 10 µg/ml human fibrinogen in a humid chamber overnight at 4°C, blocked with 1% BS in PBS for 1 h at RT and washed with Tyrode´s buffer with $Ca^{2+}$ on the following day. Platelets were washed and adjusted to a count of 300,000 platelets/µl. 30 µl platelets were resuspended in 70 µl $Ca^{2+}$ Tyrode´s buffer, stimulated with

0.01 U/ml thrombin and directly added onto the fibrinogen-coated coverslips. After adhesion for 5, 15 and 30 min, the platelets were fixed with 4% PFA in PHEM containing 0.1% TritonX100 (for following confocal microscopy) or 4% PFA in PBS (for differential interference contrast (DIC) microscopy only) for 5 min.

**Direct Stochastic Optical Reconstruction Microscopy (*d*STORM)**

Platelets were washed and adjusted to 300,000 platelets/µl in Tyrode´s buffer without $Ca^{2+}$. An 8-well chamber (Cellvis) was coated with 200 µl of human fibrinogen (10 µg/ml) for 2 h at 37°C before 60 µl platelet solution was added together with 200 µl Tyrode´s buffer with $Ca^{2+}$ and stimulated with 0.01 U/ml thrombin. After 30 min incubation at 37°C, the supernatant was removed and spread platelets were fixed in 200 µl prewarmed cytoskeleton buffer I for 2 min at 37°C, following an 18 min incubation with 200 µl cytoskeleton buffer II at RT. The samples were quenched with 0.1% NaBH4 for 15 min and blocked with 5% BSA for 1 h. For visualization of the platelet cytoskeleton, the samples were stained with either 5 µl AF647-conjugated Phalloidin (#A22287) or a total of 1 µg AF647-conjugated anti-α-tubulin antibody at 4°C overnight. One color *d*STORM samples were imaged on a widefield setup based on an inverted microscope (Olympus IX-71) equipped with an oil immersion objective (Olympus APON 60xO TIRF, NA 1.49). *d*STORM images were reconstructed using the open source software rapidSTORM 3.3.[2]

**TEM analysis of resting platelets**

Anesthetized mice were retro-orbitally bled up to 50 µl into 300 µl heparin (20 U/ml in TBS) and platelets were isolated from whole blood as previously described.[24] After fixation overnight at 4°C, the platelets were washed for three times in cacodylate buffer and centrifuged at 1500 rpm for 5 min. Subsequently, the samples were treated with 1% $OsO_4$ for 1 h at RT, washed again with cacodylate buffer, followed by addition of $H_2$Obidest and 2% uranyl acetate (in $H_2O$) for 1 h at 4°C The samples were dehydrated by an ethanol series, 10 min incubation in propylenoxide twice and 1h rotation in a 1:1 mixture of propylenoxide and epon. The propylenoxide was discarded and the samples incubated overnight at RT in epon, which was then hardened for 48 h. Ultra-thin sections were generated and stained using 2% uranyl acetate and lead citrate. Images were acquired at a JEM-2100 (JEOL). Granule and MT coil abundance was determined manually using ImageJ Software.

**TEM analysis of BM MKs**

Femora of sacrificed mice were isolated and cut into approximately 4 mm pieces and fixed Karnovsky fixative (2% PFA, 2.5% glutaraldehyde in 0.1 M cacodylate buffer) at 4°C o/n under rotation. Bones were subjected with 2% osmium tetroxide in 50 mM cacodylate buffer (pH 7.2) and subsequently stained 0.5% aqueous uranyl acetate before dehydration in an ethanol series and embedding in epon. Ultra thin sections were imaged at a JEM-2100 (JEOL).

**Hematoxylin and eosin staining on paraffin sections**

Spleens of sacrificed mice were dissected and fixed in 4% PFA at 4°C o/n and dehydrated before paraffin embedding. Sections of the organs were cut at a Leica microtome, rehydrated with an ethanol gradient, and stained with hematoxylin and eosin (H&E). The number of MKs and the general appearance of the organs were determined at a Leica DMI 4000B microscope.

**Immunofluorescence microscopy of BM cryosections**

Immunofluorescence stainings were performed as described previously.[15] Femora of sacrificed mice were isolated and fixed in phosphate-buffered saline (PBS) (Sigma-Aldrich), containing 4% PFA, o/n at 4°C and transferred to a serial gradient of sucrose in PBS solutions of 10, 20, 30% (w/v), each for 24h at 4°C. Subsequently, the bones were embedded in cryo mold dishes with SCEM medium and deep frozen at -80°C. For IF-staining bones were sliced into 7 µm sections on Kawamoto adhesive films[3] with a cryotom (Leica CM1900). Sections were thawed and rehydrated for 20 min with PBS and directly fixed for 10 min with 4% PFA in PHEM-buffer (100 mM PIPES, 5.25 mM HEPES, 10 mMEGTA, 20 mM MgCl2, pH 6.8). Sections were washed three times with a washing buffer consisting of 5% fetal calve serum (FCS) and 0.1% Tween20/ or Triton X-100 in PBS, and subsequently unspecific binding sites were blocked by using a blocking buffer, which additionally to the FCS and Tween also contained 5% goat serum (#5425, CST). To visualize platelets and MKs, sections were incubated for 45 min at RT with Alexa488-conjugated anti-GPIX antibodies (2mg/ml, Xia.B4, Emfret Analytics) and unconjugated, or Alexa647-conjugated anti-CD105 antibodies (3.33 mg/ml, MJ7/18, Biolegend), were used to stain the endothelium. After washing the slides three times, they were mounted with Fluoroshield, containing DAPI (Sigma-Aldrich), to additionally stain the nuclei. Images were taken using a TCS SP8 confocal laser scanning microscope (Leica Microsystems CMS) with a 40x oil objective 8NA 1.3). Image documentation and analysis was performed with LAS X software, for MK measurements, images were processed with FIJI software.

**MK isolation with lineage depletion**

Cells were obtained as described previously,[27] by using an antibody mixture (Biolegend) in combination with magnetic beads for lineage depletion. The antibody negative fraction was cultured in DMEM (Gibco) containing 50 ng/ml TPO and 100 U/ml rHirudin for 72h. Afterwards, the cells were enriched with the help of BSA density gradient and used to perform experiments.

**Spreading of BM-derived *in vitro*-differentiated MKs on matrices**

Glass coverslips were coated with HORM collagen (50 µg/ml) for 3h at 37°C or o/n at 4°C and washed with PBS once. 500 µl MK cell suspension were pipetted onto the slides and MKs were allowed to adhere to the coverslips for 3h at 37°C and 5% $CO_2$. The cells were then fixed and permeabilized in PHEM fixation buffer for 20 min and blocked with 3% BSA/PBS for 30 min before staining with fluorescently labeled antibodies o/n at 4°C. Afterwards, MKs were washed twice with PBS and once with $H_2O$ and were mounted using Fluoroshield. Images were taken using a TCS SP8 confocal laser scanning microscope (Leica Microsystems CMS) with a 40x oil objective 8NA 1.3). Image documentation and analysis was performed with LAS X software, for MK measurements, images were processed with FIJI software

**Immunoblotting of platelet and MK lysates**

Washed platelets adjusted to a concentration of $1x10^6$/µl were lysed with IP-buffer (15 mM TRIS HCl, 155 mM NaCl, 1mM EDTA. 0.005% $NaN_3$), supplemented with 2% NP-40 and 1x protease inhibitors for 20 min on ice. For immunoblotting, samples were mixed with 4x reducing Laemmli buffer and boiled for 5 min at 95°C. For MK lysates, BM-derived *in vitro*-differentiated MKs were enriched,[27] and were collected in protein low binding tubes (Eppendorf) and washed with 1 ml PBS containing 3 mM EDTA. According to the pellet size, cells were immediately lysed in an appropriate volume of 1x RIPA, buffer supplemented with 1x Halt protease and phosphatase inhibitors for 20 min on ice and centrifuged for 10 min at max rpm. The supernatant was collected and the protein content determined by a *Bicinchoninic acid assay* (BCA) protein assay. Samples were adjusted with 4x reducing sample buffer and boiled for 5 min at 95°C before immunoblotting. Platelet and MK lysates were analyzed by SDS-PAGE and immunoblotting using polyvinylidene difluoride (PVDF) membranes, which were incubated with the indicated antibodies, before visualization using an Amersham Image 680 (GE Healthcare).

**SUPPLEMENTAL TABLES**

| | | *wt* | | *RhoB*$^{-/-}$ | | |
|---|---|---|---|---|---|---|
| | | **mean** | **SD** | **mean** | **SD** | **P-value** |
| WBC# | $10^3/mm^3$ | 7.2 | 4.0 | 7.4 | 1.9 | 0.89 |
| LYM# | $10^3/mm^3$ | 4.9 | 2.5 | 5.6 | 1.5 | 0.53 |
| MON# | $10^3/mm^3$ | 0.3 | 0.3 | 0.3 | 0.1 | 0.61 |
| GRA# | $10^3/mm^3$ | 1.9 | 1.4 | 1.5 | 0.6 | 0.46 |
| EOS# | $10^3/mm^3$ | 0.0 | 0.1 | 0.0 | 0.0 | 0.80 |
| LYM% | % | 72.0 | 6.5 | 76.6 | 6.4 | 0.18 |
| MON% | % | 4.9 | 1.0 | 4.6 | 0.8 | 0.47 |
| GRA% | % | 23.1 | 5.7 | 18.8 | 5.8 | 0.16 |
| EOS% | % | 1.2 | 1.4 | 08 | 0.3 | 0.49 |
| RBC | $10^6/mm^3$ | 8.6 | 0.4 | 8.0 | 0.5 | 0.02 |
| HGB | g/dl | 15.4 | 0.9 | 15.1 | 1.1 | 0.48 |
| HCT | % | 49.3 | 3.2 | 47.9 | 4.7 | 0.53 |
| MCV | $\mu m^3$ | 57.3 | 3.3 | 60.0 | 2.8 | 0.10 |
| MCH | pg | 18.0 | 0.7 | 18.8 | 0.5 | 0.01 |
| MCHC | g/dl | 31.3 | 0.8 | 31.5 | 0.9 | 0.73 |
| RDW | % | 14.7 | 1.6 | 13.7 | 0.2 | 0.10 |

**Supplemental Table 1. Peripheral blood cell counts of *wt* and *RhoB*$^{-/-}$ mice.** Blood parameters were measured using a scil Vet abc Plus+ blood analyzer (n= at least 7 mice per genotype). Mann-Whitney test, mean ± SD.

| | *wt* | | *RhoB*$^{-/-}$ | | |
|---|---|---|---|---|---|
| | **Mean** | **SD** | **Mean** | **SD** | **P-value** |
| GPIb | 277 | 29 | 259 | 33 | 0.43 |
| GPV | 155 | 2 | 135 | 3 | <0.01 |
| GPIX | 339 | 44 | 317 | 55 | 0.26 |
| GPVI | 39 | 8 | 32 | 4 | 0.01 |
| CLEC-2 | 109 | 38 | 125 | 35 | 0.30 |
| CD9 | 970 | 184 | 948 | 234 | 0.79 |
| α2 | 37 | 7 | 38 | 8 | 0.86 |
| β1 | 108 | 11 | 108 | 4 | 0.97 |
| aIIbβ3 | 403 | 70 | 407 | 122 | 0.93 |

**Supplemental Table 2. Surface expression of glycoproteins of resting platelets.** Levels of the main glycoproteins on *wt* and *RhoB*$^{-/-}$ platelets were analyzed by flow cytometry using FITC-labeled antibodies (n=4 mice per genotype). Mann-Whitney test, mean ± SD.

#### SUPPLEMENTAL FIGURE LEGENDS

**Supplemental Figure 1. Normal granule count and integrin β1 activation in *RhoB*$^{-/-}$ platelets.** (A-B) Number of α and dense (δ-) granules was determined by manual counting of twenty TEM images of resting *wt* and *RhoB*$^{-/-}$ platelets/mouse using 3 mice/genotype. Each data point represents the granule count of one single platelet (n=60). (C) Platelet integrin β1 activation was analyzed by flow cytometry using 9EG7-FITC binding upon stimulation with classical agonists at different concentrations (n=4). Mann-Whitney test, mean ± SD. Rest = resting; ADP/U466 = Adenosine diphosphate/U46619; Thr = thrombin; CRP = collagen-related peptide; CVX = convulxin; Rhd = Rhodocytin.

**Supplemental Figure 2. Unaltered F-actin and MT distribution in *RhoB*$^{-/-}$ MKs upon adhesion on collagen *in vitro*.** (A) Representative confocal images of BM-derived *in vitro*-differentiated MKs 24 h after BSA gradient. The MKs were spun down to Poly L-Lysine coated glass slides and stained with phalloidin-Atto647 (magenta), α-tubulin-Alexa488 (cyan) and DAPI to visualize the nucleus. White arrowheads mark proplatelet tips and shafts. Scale bar in overview: 40 µm; inset: 5 µm. (B) Representative confocal microscopy images of BM-derived *in vitro*-differentiated MKs spread on collagen. MKs were enriched by BSA gradient and allowed to spread on HORM collagen-coated glass slides for 3 h, formaldehyde fixed, blocked and stained with phalloidin-Atto647 (magenta), α-tubulin-Alexa488 (cyan) and DAPI to stain the nucleus. Scale bar in overview; 40 µm; inset: 5 µm. (C) Quantification of MK podosome density on images described in (B), normalized to *wt*, pooled data of 3 independent experiments, 3 mice/genotype. Each data point represents one MK (*wt* = 26; *RhoB*$^{-/-}$ = 28). (D) Quantification of the MK spreading area on images described in (B), normalized to *wt*, pooled data of 3 independent experiments, 3 mice/genotype. At least 5 images/mouse were analyzed. Each data point represents one MK (*wt* = 56; *RhoB*$^{-/-}$ = 82). (E) Quantification of MK F-actin content on images described in (B), normalized to *wt*, pooled data of 3 independent experiments, 3 mice/genotype. At least 5 images/mouse were analyzed. Each data point represents one MK (*wt* = 41; *RhoB*$^{-/-}$ = 56). (F) Quantification of α-tubulin fluorescent intensity on images described in (B), normalized to *wt*, pooled data of 2 independent experiments, 3 mice/genotype. At least 5 images/mouse were analyzed. Each data point represents one MK (*wt* = 28; *RhoB*$^{-/-}$ = 27). Mann-Whitney test, mean ± SD. AU = arbitrary unit; MFI = mean fluorescence intensity.

**Supplemental Figure 3. Actin and tubulin cytoskeleton components and regulators in *RhoB*$^{-/-}$ MKs.** (A-C) Representative blots of BM-derived *in vitro*-differentiated MK lysates from *wt* and *RhoB*$^{-/-}$ mice immunoblotted for the indicated proteins or PTMs. GAPDH was used as loading control (n=3). (D) Analysis of F-actin assembly upon platelet activation. Left: MFI of phalloidin-FITC in resting platelets and platelets stimulated with the indicated agonists. Right:

Ratio of CRP or thrombin activated and resting values (n=3). Mann-Whitney test, mean ± SD. (E) Representative confocal immunofluorescence microscopy images of MT organization after disassembly at 4°C following reassembly at 37°C in *wt* and *RhoB*$^{-/-}$ platelets. Platelets were stained with phalloidin-Atto647 (magenta) and α-tubulin-Alexa488 (cyan). Scale bar in overview: 10 μm.

### Supplemental Figure 1

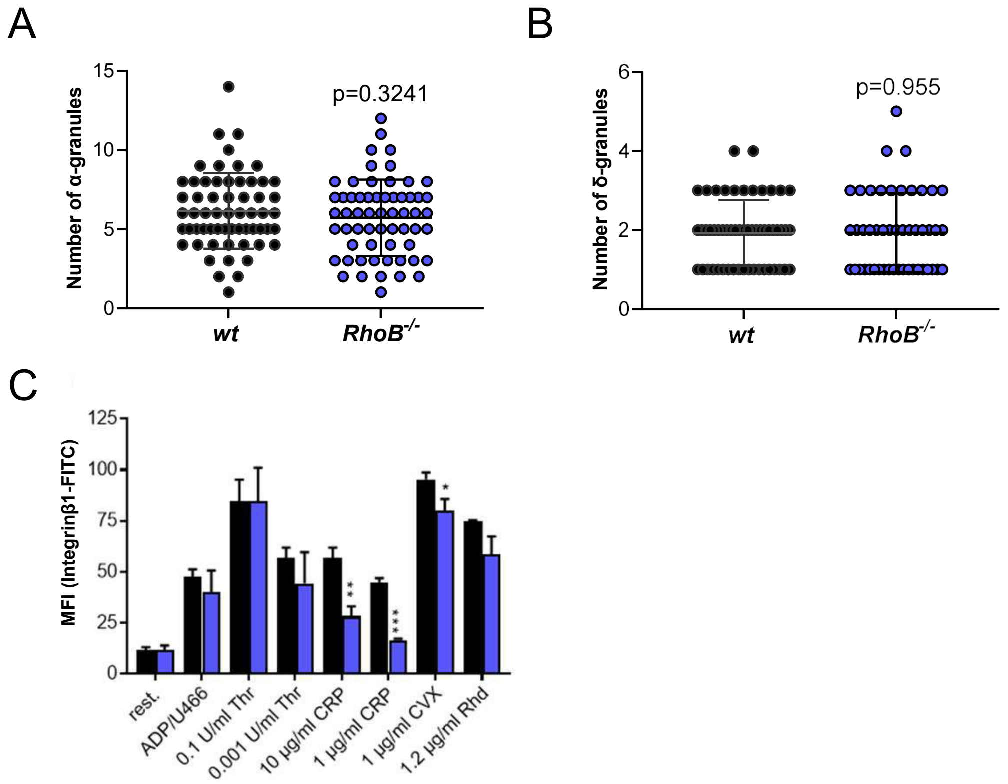

### Supplemental Figure 2

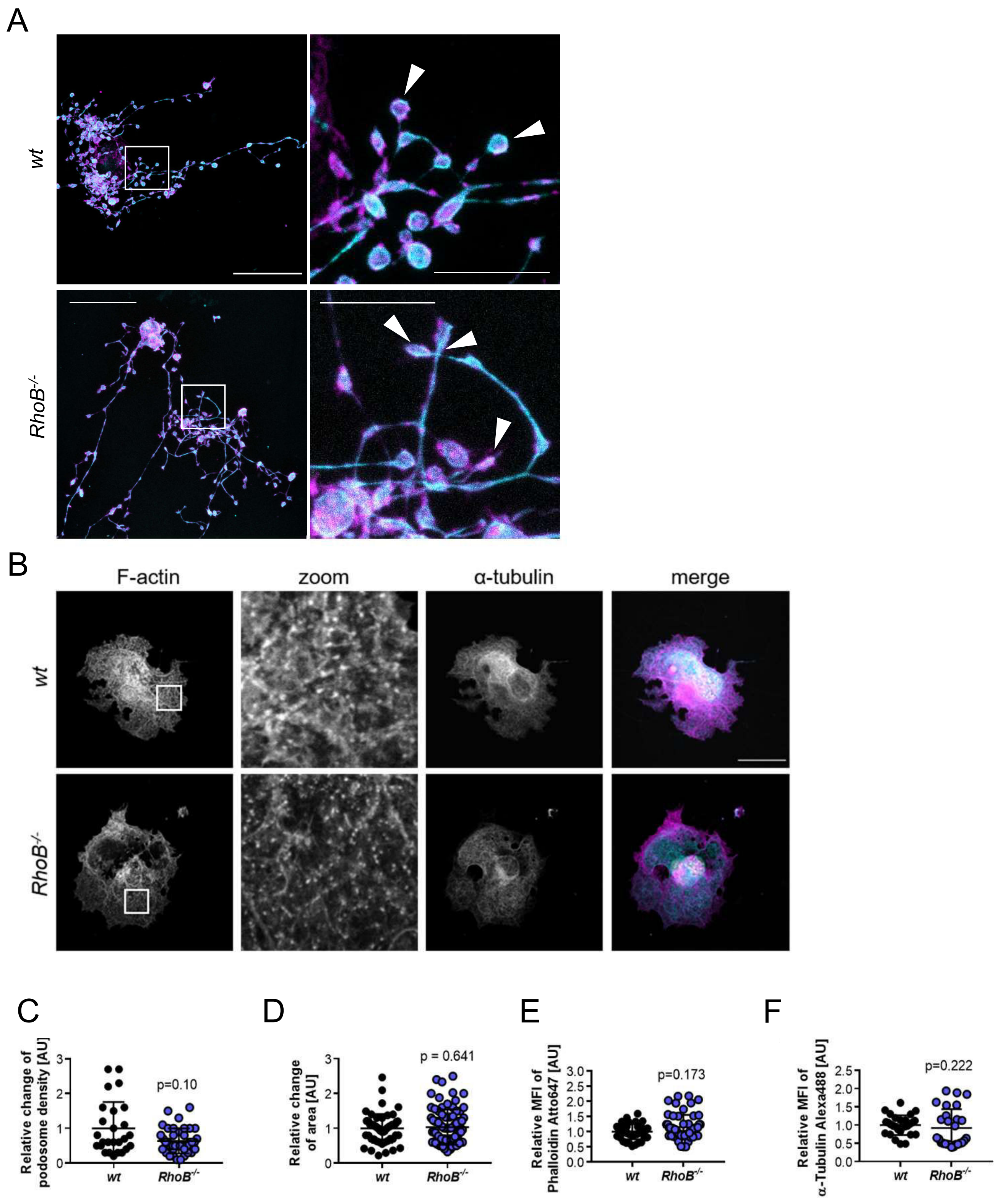

### Supplemental Figure 3

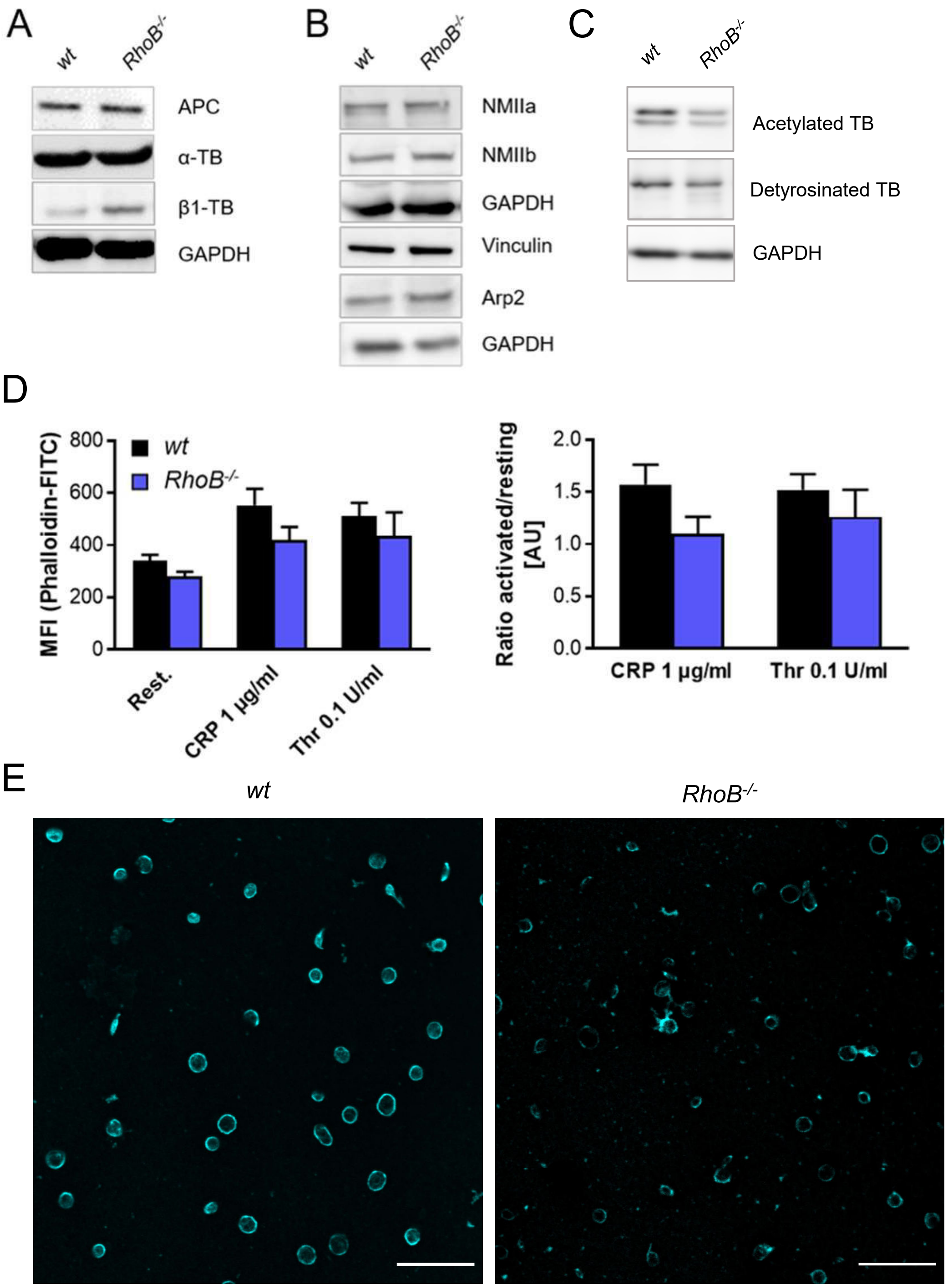
